# Decoding Motor Plans Using a Closed-Loop Ultrasonic Brain-Machine Interface

**DOI:** 10.1101/2022.11.10.515371

**Authors:** Whitney S. Griggs, Sumner L. Norman, Thomas Deffieux, Florian Segura, Bruno-Félix Osmanski, Geeling Chau, Vasileios Christopoulos, Charles Liu, Mickael Tanter, Mikhail G. Shapiro, Richard A. Andersen

**Affiliations:** Biology and Biological Engineering, California Institute of Technology, Pasadena, CA 91125, USA; Physics for Medicine Paris, INSERM, CNRS, ESPCI Paris, PSL Research University, 75012 Paris, France; INSERM Technology Research Accelerator in Biomedical Ultrasound, Paris, France; Iconeus, 6 rue Jean Calvin, Paris, France; T&C Chen Brain-Machine Interface Center, California Institute of Technology, Pasadena, CA 91125, USA; Bioengineering, University of California, Riverside, Riverside, CA, USA; Department of Neurological Surgery, Keck School of Medicine of USC, Los Angeles, CA 90033, USA; USC Neurorestoration Center, Keck School of Medicine of USC, Los Angeles, CA 90033, USA; Rancho Los Amigos National Rehabilitation Center, Downey, CA 90242, USA; Chemistry & Chemical Engineering, California Institute of Technology, Pasadena, CA 91125, USA; Howard Hughes Medical Institute, Pasadena, CA 91125, USA

## Abstract

Brain-machine interfaces (BMIs) can be transformative for people living with chronic paralysis. BMIs translate brain signals into computer commands, bypassing neurological impairments and enabling people with neurological injury or disease to control computers, robots, and more with nothing but thought. State-of-the-art BMIs have already made this future a reality in limited clinical trials. However, high performance BMIs currently require highly invasive electrodes in the brain. Device degradation limits longevity to about 5 years. Their field of view is small, restricting the number, and type, of applications possible. The next generation of BMI technology should include being longer lasting, less invasive, and scalable to sense activity from large regions of the brain. Functional ultrasound neuroimaging is a recently developed technique that meets these criteria. In this present study, we demonstrate the first online, closed-loop ultrasonic brain-machine interface. We used 2 Hz real-time functional ultrasound to measure the neurovascular activity of the posterior parietal cortex in two nonhuman primates (NHPs) as they performed memory-guided movements. We streamed neural signals into a classifier to predict the intended movement direction. These predictions controlled a behavioral task in real-time while the NHP did not produce overt movements. Both NHPs quickly succeeded in controlling up to eight independent directions using the BMI. Furthermore, we present a simple method to “pretrain” the BMI using data from previous sessions. This enables the BMI to work immediately from the start of a session without acquiring extensive additional training data. This work establishes, for the first time, the feasibility of an ultrasonic BMI and prepares for future work on a next generation of minimally invasive BMIs that can restore function to patients with neurological, physical, or even psychiatric impairments.

## MAIN

Brain-machine interfaces (BMIs) translate complex brain signals into computer commands and are a promising method to restore the capabilities of human patients with paralysis^1^. State-of-the-art BMIs have already made this future a reality in limited clinical trials^2–6^. However, these BMIs require invasive electrode arrays inserted into the brain. Device degradation limits the BMI’s longevity to typically around 5 years^7,8^. The implants only sample from small regions of superficial cortex. These are some of the factors limiting BMI technology’s adoption to a broader patient population.

Functional ultrasound (fUS) imaging is a recently developed technique that is poised to enable longer lasting, less invasive BMIs that can scale to sense activity from large regions of the brain. fUS neuroimaging uses ultrafast pulse-echo imaging to sense changes in cerebral blood volume (CBV)^9^. It has a high sensitivity to slow blood flow (∼1mm/s velocity) and balances good spatiotemporal resolution (100 μm; <1 sec) with a large and deep field of view (∼7 centimeters). Previously, we demonstrated that fUS neuroimaging possesses the sensitivity and field of view to decode movement intention on a single-trial basis simultaneously for two directions (left/right), two effectors (hand/eye), and task state (go/no-go)^10^. However, we performed this *post hoc* (off-line) analysis using pre-recorded data. In this study, we demonstrate the first online, closed-loop, functional ultrasound brain-machine interface (fUS-BMI). In addition, we present key advances that build on previous fUS neuroimaging studies. These include decoding eight movement directions and designing decoders stable across more than 40 days.

## RESULTS

We used a miniaturized 15.6-MHz, 128-channel, linear ultrasound transducer array paired with a realtime ultrafast ultrasound acquisition system to stream 2 Hz fUS images from nonhuman primates (NHPs) as they performed memory-guided eye movements (**Fig. 1**). We positioned the transducer surface normal to the brain above the dura mater and recorded from coronal planes of the left posterior parietal cortex (PPC), a sensorimotor association area that uses multisensory information to guide movements and attention^11–13^. This technique achieved a large field of view (12.8 mm width, 16 mm depth, 400 μm plane thickness) while maintaining high spatial resolution (100 μm × 100 μm in-plane). This allowed us to stream high-resolution hemodynamic changes across multiple PPC regions simultaneously, including the lateral (LIP) and medial (MIP) intraparietal cortex (**Fig. 1A**). Previous research has shown that the LIP and MIP are involved in planning eye and reach movements respectively^14–16^. This makes PPC a good region from which to record effector-specific movement signals.

**Figure 1:**
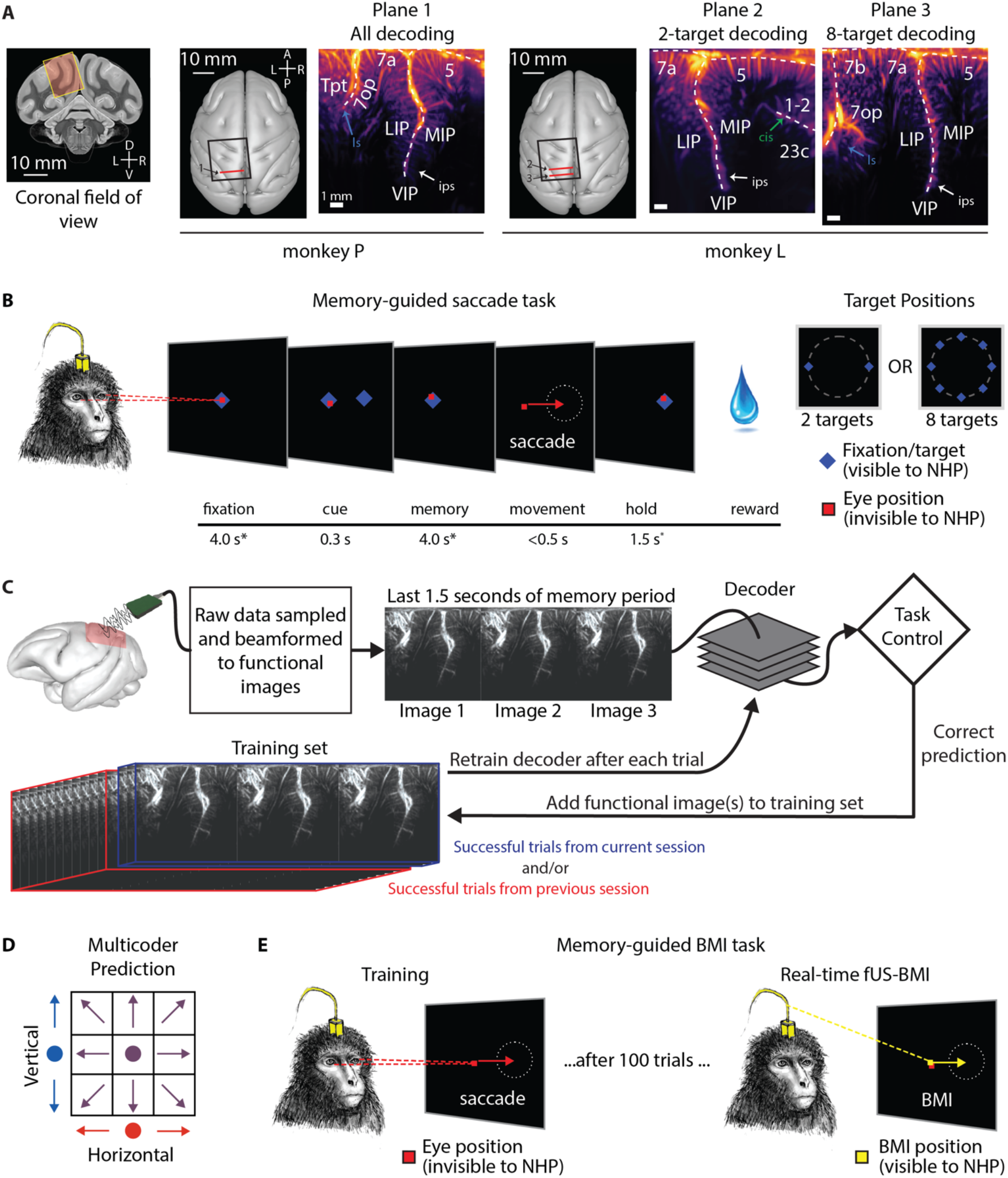
Anatomical recording planes and behavioral tasks. **(A)** Coronal fUS imaging planes used for monkey P and monkey L. A coronal slice from an MRI atlas shows the approximate field of view for the fUS imaging plane. The 24 × 24 mm (inner dimension) chambers were placed surface normal to the skull above a craniotomy (black square). The ultrasound transducer was positioned to acquire a consistent coronal plane across different sessions (red line). The vascular maps show the mean Power Doppler image from a single imaging session. The three imaging planes were chosen for good decoding performance in a pilot offline dataset. Anatomical labels based upon^17^. **(B)** Memory-guided saccade task. A trial started with the NHP fixating on a center fixation cue. A target cue was flashed in a peripheral location. During the memory period, the NHP continued to fixate on the center cue and planned an eye movement to the peripheral target location. Once the fixation cue was extinguished, the NHP performed a saccade to the remembered location and maintained fixation on the peripheral location before receiving a liquid reward. *The fixation, memory, and hold periods were subject to ±500 ms of jitter (uniform distribution) to prevent the NHP from anticipating task state changes. The peripheral cue was chosen from two or eight possible target locations depending on the specific experiment. Red square represents the NHP’s eye position but is not visible to the NHP. **(C)** fUS-BMI algorithm. Real-time 2 Hz functional images were streamed to a linear decoder that controlled the behavioral task. The decoder used the last 3 fUS images (1.5 seconds) of the memory period to make its prediction. If the prediction was correct, the data from that prediction was added to the training set. The decoder was retrained after every successful trial. The training set consisted of trials from the current session and/or from a previous fUS-BMI session. **(D)** Multicoder algorithm. For predicting eight movement directions, the vertical component (blue; up, middle, or down) and the horizontal component (red; left, middle, or right) were separately predicted and then combined to form each fUS-BMI prediction (purple). **(E)** Memory-guided BMI task. The BMI task is the same as (B) except that the movement period is controlled by the brain activity (via fUS-BMI) rather than eye movements. After 100 successful eye-movement trials, the fUS-BMI controlled the movement prediction (closed-loop control). During the closed-loop mode, the NHP had to maintain fixation on the center fixation cue until reward delivery. If the BMI prediction was correct and the NHP held fixation correctly, the NHP received a large liquid reward (1 sec; ∼0.75 mL). If the BMI prediction was incorrect, but the NHP held fixation correctly, the NHP received a smaller liquid reward (100 ms; ∼0.03 mL). Red square represents the NHP’s eye position and is not visible to the monkey. Yellow square represents the BMI-controlled cursor position and is visible to the monkey.

We streamed the real-time 2 Hz fUS images into a BMI decoder that used principal component (PCA) and linear discriminant analysis (LDA) to predict planned movement directions. We used the BMI output to directly control the behavioral task (**Fig. 1C**). To build the initial training set for the decoder, each of two NHPs initially performed instructed eye movements to a randomized set of two or eight peripheral targets. We used the fUS activity during the delay period preceding successful eye movements to train the decoder. After 100 successful training trials, we switched to closed-loop BMI mode where the movement came from the fUS-BMI (**Fig. 1E**). During this closed-loop BMI mode, the NHP continued to fixate on the center cue until after the delivery of the liquid reward. During the interval between a successful trial and the subsequent trial, we rapidly retrained the decoder, continuously updating the decoder model as each NHP used the fUS-BMI.

### Online decoding of two eye movement directions

To demonstrate the feasibility of a fUS-BMI, we first performed online, closed-loop decoding of two movement directions (**Fig. 2**). After building a preliminary training set of 20 trials, we began testing the decoder’s accuracy on each new trial in a training mode not visible to the NHP (blue line, **Fig. 2A**). After 100 trials, we switched the BMI from training to closed-loop decoding where the NHP now controlled the task direction using his movement intention, i.e., the brain activity detected by the fUS-BMI in the last 3 fUS images of the memory period (**Fig. 1C, E, Fig. 2A-yellow line**). At the conclusion of each trial, the NHP received visual feedback of the fUS-BMI prediction. In our second closed-loop 2-direction session, the decoder reached significant accuracy (p<0.01; 1-sided binomial test) after 55 training trials and improved in accuracy until peaking at 82% accuracy at trial 114 (**Fig 2A**). The decoder predicted both directions well above chance level but displayed better performance for rightward movements (**Fig. 2B**). To understand which brain regions were most important for the decoder performance, we performed a searchlight analysis with a 200 μm, i.e., 2 voxel, radius (**Fig. 2C**). Dorsal LIP and Area 7a contained the voxels most informative for decoding intended movement direction.

**Figure 2:**
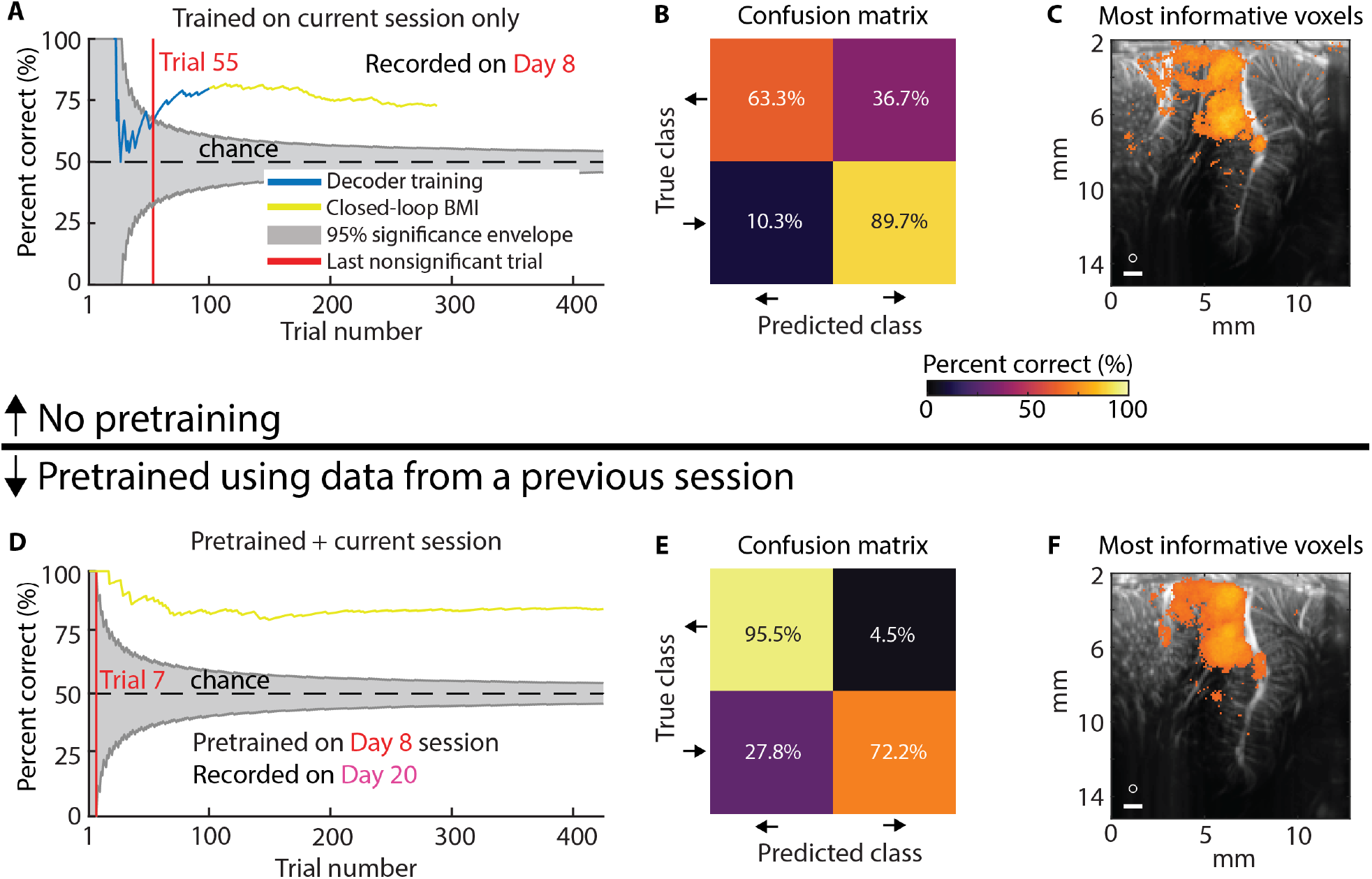
Example sessions decoding two saccade directions. **(A)** Mean decoding accuracy as a function of trial number. Blue represents fUS-BMI training where the NHP controlled the task using overt eye movements. The BMI performance shown in blue was generated post hoc with no impact on the real-time behavior. Yellow represents trials under fUS-BMI control where the NHP maintained fixation on the center cue and the task direction was controlled by the fUS-BMI. Grey chance envelope – 95% binomial distribution. Red line – last nonsignificant trial. **(B)** Confusion matrix of decoding accuracy across entire session represented as percentage (rows add to 100%). **(C)** Searchlight analysis represents the 10% voxels with the highest decoding accuracy (threshold is q<=1.66e-6). White circle – 200 μm searchlight radius used. White bar – 1 mm scale bar. **(D-F)** Same format as in (A-C). fUS-BMI was pretrained using data collected from a previous session. Top 10% of voxels in searchlight analysis corresponds to a threshold of q<=2.07e-13. Text color for the day labels matches the day label colors in **Fig. 3**.

An ideal BMI needs very little training data and no retraining between sessions. Electrode-based BMIs typically require calibration or re-training for each subsequent session, due to difficulty in recording from the same neurons across multiple days^18^.Thanks to its wide field of view, fUS neuroimaging can image from the same brain regions over time, and therefore may be an ideal technique for stable decoding across many sessions. To test this hypothesis, we pretrained the fUS-BMI using a previously recorded session’s data and then tested the decoder in an online experiment. To perform this pretraining, we first aligned the data from the previous session’s imaging plane to the current session’s imaging plane (**Fig. S1**). We used semi-automated rigid body registration to find the transform between the previous and current imaging plane, applied this 2D image transform to each frame of the previous session, and saved the aligned data. This semi-automated pre-registration process took less than 1 minute. To pretrain the model, the fUS-BMI automatically loaded this aligned dataset and trained the initial decoder. The fUS-BMI reached significant performance substantially faster (**Fig. 2D**) when we used this pretraining approach. The fUS-BMI achieved significant accuracy at Trial 7, approximately 15 minutes faster than the example session without pretraining.

To quantify the benefits of pretraining upon fUS-BMI training time and performance, we compared fUS-BMI performance across all sessions when (a) using only data from the current session versus (b) pretraining with data from a previous session (**Fig. 3**). For all real-time sessions that used pretraining, we also created a *post hoc* (offline) simulation of the fUS-BMI results without using pretraining. For each simulated session, we passed the recorded data through the same classification algorithm used for the real-time fUS-BMI but did not use any data from a previous session.

**Figure 3.**
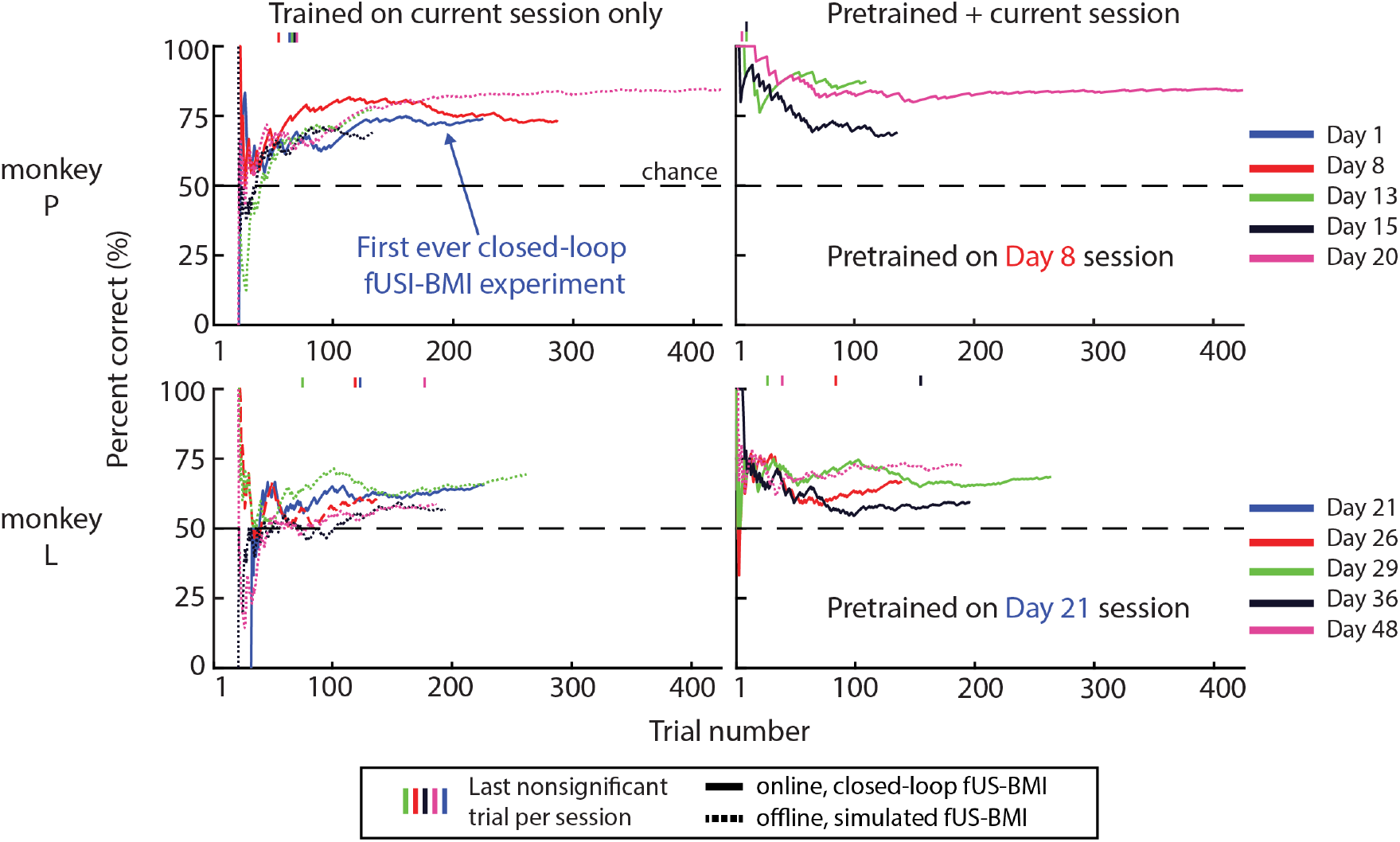
Performance across sessions for decoding two saccade directions. Mean decoder accuracy during each session for monkey P and L. Solid lines are real-time results while fine-dashed lines are simulated sessions from post hoc analysis of real-time fUS imaging data. Vertical marks above each plot represent the last nonsignificant trial for each session. Day number is relative to the first fUS-BMI experiment. Coarse-dashed horizontal black line represents chance performance.

#### Using only data from the current session

The mean decoding accuracy reached significance (p<0.01; 1-sided binomial test) at the end of each online, closed-loop recording session (2/2 sessions monkey P, 1/1 session monkey L) and most offline, simulated recording sessions (3/3 sessions monkey P, 3/4 sessions monkey L) (**Fig. 3**). For monkey P, decoder accuracies reached 75.43 ± 2.56% correct (mean ± SEM) and took 40.20 ± 2.76 trials to reach significance. For monkey L, decoder accuracies reached 62.30 ± 2.32% correct and took 103.40 ± 23.63 trials to reach significance.

#### Pretraining with data from a previous session

The mean decoding accuracy reached significance at the end of each online, closed-loop recording session (3/3 sessions monkey P, 4/4 session monkey L) (**Fig. 2C**). Using previous data reduced the time to achieve significant performance (100% of sessions reached significance sooner, monkey P – 36-43 trials faster; monkey L – 15-118 trials faster). The performance at the end of the session was not statistically different from performance in the same sessions without pretraining (paired t-test, p<0.05). For monkey P, accuracies reached 80.21 ± 5.61% correct and took 9.00 ± 1 trials to reach significance. For monkey L, accuracies reached 66.78 ± 2.79% correct and took 71.00 ± 28.93 trials to reach significance. Assuming no missed trials, pre-training decoders shortened training by 10-45 minutes. We also simulated the effects of not using any training data from the current session, i.e., using only the pretrained model (**Fig. S2A**). We did not observe a statistically significant difference between the performance (accuracy or number of trials to significant performance) for either NHP whether current session training data was included or not.

These results demonstrate online decoding of fUS signals into two directions of movement intention, NHPs learning to control the task using the fUS-BMI, and that pretraining using a previous session’s data greatly reduced, or even eliminated, the amount of new training data required in a new session.

### Online decoding of eight eye movement directions

Having demonstrated that we could achieve similar, **but online and closed-loop**, performance to our previous offline decoding paper^10^, we extended the capabilities of fUS-BMI by decoding eight movement directions in real time (**Fig. 4**). We used a “multicoder” architecture where we predicted the vertical (up, middle, or down) and horizontal (left, middle, or right) components of intended movement separately and then combined those independent predictions to form a final prediction (e.g., up and to the right) (**Fig. 1D**). In the first 8-direction experiment, the decoder reached significant accuracy (p<0.01; 1-sided binomial test) after 86 training trials and improved until plateauing at 34-37% accuracy (**Fig 4A** – upper plot), compared to 12.5% chance level, with most errors indicating directions neighboring the cued direction (**Fig. 4B**). To capture this extra information about proximity of each prediction to the true direction, we examined the mean absolute angular error. The fUS-BMI reached significance at 55 trials and steadily decreased its mean absolute angular error to 45° by the end of the session (**Fig 4A** – bottom plot). Compared to the most informative voxels for the 2-target eye decoder, a larger portion of LIP, including ventral LIP, contained the most informative voxels for decoding eight directions of movement (**Fig. 4C**).

**Figure 4.**
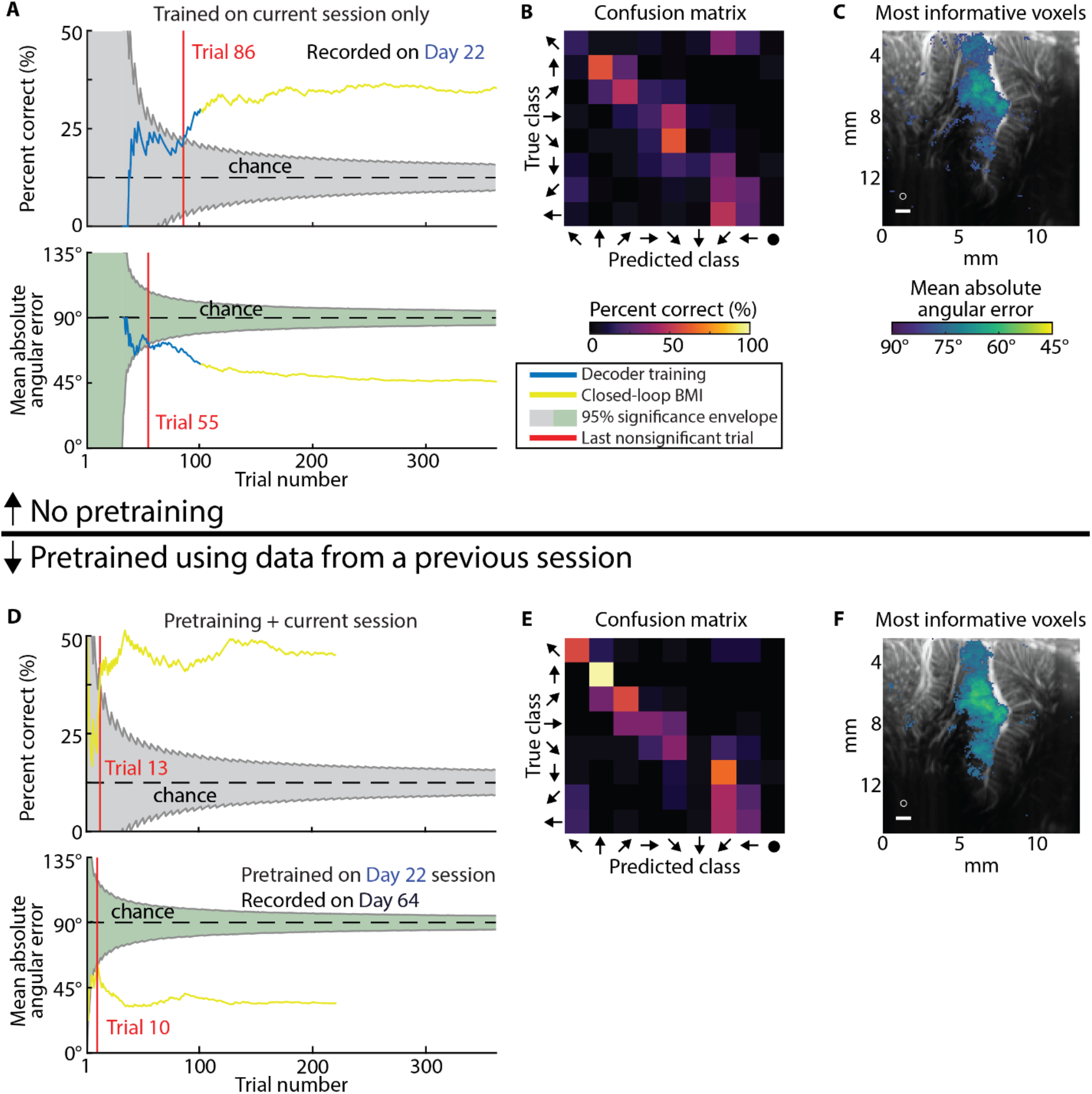
Example sessions decoding eight saccade directions. **(A)** Mean decoding accuracy and mean absolute angular error as a function of trial number. Blue represents fUS-BMI training where the NHP controlled the task using overt eye movements. The BMI performance shown here was generated post hoc tested on each new trial with no impact on the real-time behavior. Yellow represents trials under fUS-BMI control where the NHP maintained fixation on the center cue and the movement task direction was controlled by the fUS-BMI. Grey chance envelope – 95% binomial distribution. Green chance envelope – 95% permutation test distribution. Red line – last nonsignificant trial. **(B)** Confusion matrix of decoding accuracy across entire session represented as percentage (rows add to 100%). The horizontal axis plots the predicted movement (predicted class) and the vertical axis the matching directional cue (true class). **(C)** Searchlight analysis represents the 10% of voxels with the lowest mean absolute angular error (threshold is q≤2.98e-3). White circle – 200 μm searchlight radius used. White bar – 1 mm scale bar. **(D-F)** Same format as in (A-C). fUS-BMI was pretrained on data from Day 22 and updated after each successful trial. Top 10% of voxels in searchlight analysis corresponds to a threshold of q≤8e-5. Text color for the day labels matches the day label colors in **Fig. 5**.

We next tested whether pretraining would aid the 8-target decoding similarly to the 2-target decoding. As before, pretraining improved the number of trials required to reach significant decoding (**Fig. 4D**). The fUS-BMI reached significant accuracy at Trial 13, approximately 25 minutes earlier than using only data from the current session. The mean decoder accuracy reached 45% correct with a final mean absolute angular error of 34°, which was better than the performance achieved in the example session without pretraining. The searchlight analysis indicated the same regions within LIP provided the most informative voxels for decoding (**Fig. 4F**) for both the example sessions with and without pretraining. Notably, we pretrained the fUS-BMI on data from 42 days before the current session. This demonstrates that fUS-BMI can remain stable over at least 42 days. It further demonstrates that we can consistently locate the same imaging plane and that mesoscopic PPC populations consistently encode for the same directions on the time span of >1 month.

#### Using only data from the current session (**Fig. 5**-left)

The mean decoder accuracy reached significance by the end of all real-time (2/2) and simulated sessions (8/8). The mean absolute angular error for monkey P reached 45.26 ± 3.44° and the fUS-BMI took 30.75 ± 12.11 trials to reach significance. The mean absolute angular error for monkey L reached 75.06 ± 1.15° and the fUS-BMI took 132.33 ± 20.33 trials to reach significance.

**Figure 5.**
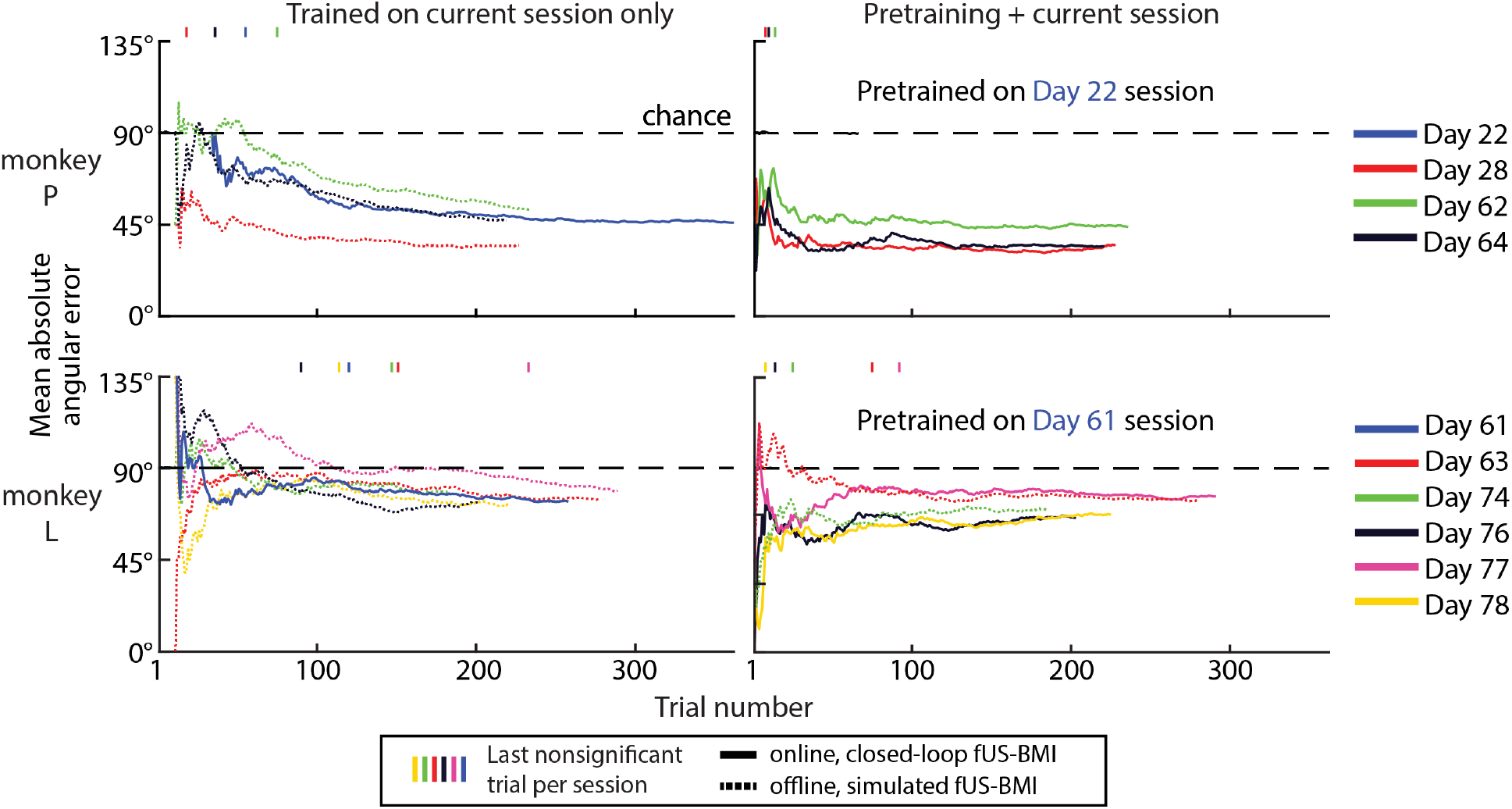
Performance across sessions for decoding eight saccade directions. Mean absolute angular error during each session for monkey P and L. Solid lines are real-time results while fine-dashed lines are simulated sessions from post hoc analyses of real-time fUS data. Vertical marks represent the last nonsignificant trial for each session. Day number is relative to the first fUS-BMI experiment. Coarse-dashed horizontal black line – chance performance.

#### Pretraining with data from a previous session (**Fig. 5**-right)

The mean decoder accuracy reached significance by the end of all real-time (6/6) and simulated sessions (2/2). The fUS-BMI reached significant decoding earlier for most sessions compared to simulated *post hoc* data; 5/5 faster monkey L; 2/3 faster monkey P (third session reached significance equally fast). For monkey P, the pretrained decoders reached significance 0-51 trials faster and for monkey L, the pretrained decoders reached significance 66-132 trials faster. For most sessions, this would shorten training by up to 45 minutes. The performance at the end of each session was not statistically different from performance in the same session without pretraining (paired t-test, p<0.05). The mean absolute angular error for monkey P reached 37.82° ± 2.86° and the fUS-BMI took 10.67 ± 1.76 trials to reach significance. The mean absolute angular error for monkey L reached 71.04° ± 2.29° and the fUS-BMI took 42.80 ± 17.05 trials to reach significance. We also simulated the effects of not using any training data from the current session, i.e., using only the pretrained model (**Fig. S2B**). We did not observe a statistically significant difference between the performance (accuracy, mean absolute angular error, or number of trials to significant performance) for either NHP whether current session training data was included or not.

These results demonstrate online decoding of fUS signals into eight directions of movement intention, a significant advance over decoding contra- and ipsilateral movements. They also show that the directional encoding within PPC mesoscopic populations is stable across at least one month, thus allowing us to reduce, or even eliminate, the need for new training data.

### Online decoding of two hand movement directions

Another strength of fUS neuroimaging is its wide field of view capable of sensing activity from multiple functionally diverse brain regions, including those that encode different movement effectors, e.g., hand and eye. To test this, we decoded intended hand movements to two target directions (reaches to the Left/Right for monkey P) in addition to our previous results decoding eye movements (**Fig 6**). The NHP performed a similar task wherein he had to maintain touch on a center dot and touch the peripheral targets during the training (**Fig. 6A**). In this scenario, we no longer constrained the monkey’s eye position, instead recording hand movements to train the fUS-BMI. After the training period, the monkey controlled the task using the fUS-BMI while keeping his hand on the center fixation cue. Notably, we used the same imaging plane used for eye movement decoding, which contained both LIP (important for eye movements) and MIP (important for reach movements). In an example session using only data from the current session (**Fig. 6B**), it took 70 trials to reach significance and achieved a mean decoder accuracy of 61.3%. The decoder predominately guessed left (**Fig. 6C**). Two foci within the dorsal LIP and scattered voxels throughout Area 7a and the temporo-parietal junction (Area Tpt) contained the most informative voxels for decoding the two movement directions (**Fig. 6D**).

**Figure 6.**
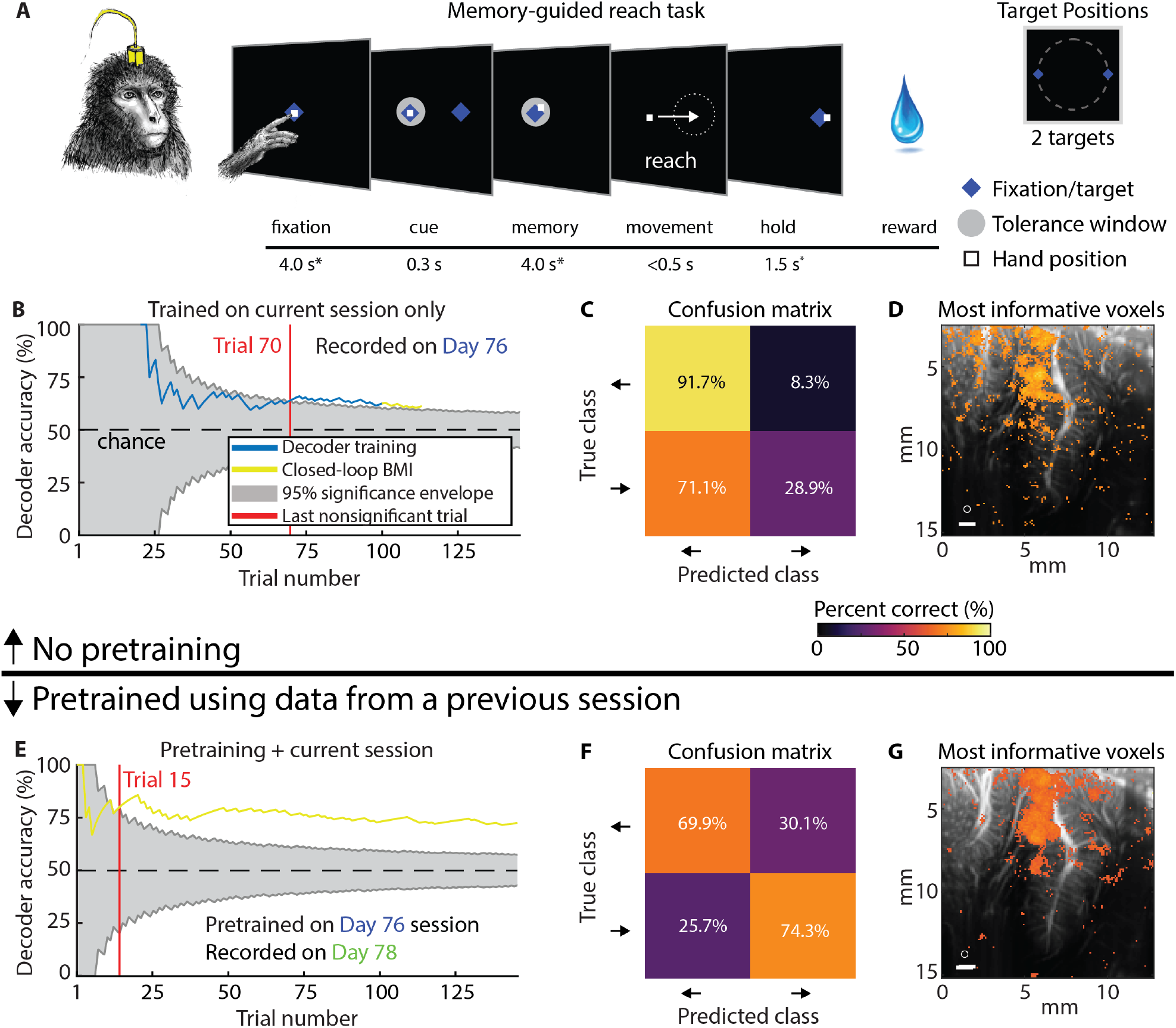
Example sessions decoding two reach directions. **(A)** Memory-guided reach task for Monkey P. Identical to memory-guided saccade task in **Fig. 1B** with all fixation or eye movements being replaced by maintaining touch on the screen and reach movements respectively. Peripheral cue was chosen from one of two peripheral targets. **(B-D)** Example results from session trained on data from the current session only. Same format as **Fig. 2A-C**. Threshold for searchlight analysis is q≤3.05e-3. **(E-G)** Example results from the Day 78 session pretrained on data from Day 76 and retrained after each successful trial. Same format as **Fig. 2D-F**. Threshold for searchlight analysis is q≤6.47e-3. Text color for the day labels matches the day label colors in **Fig. 7**.

To assess the benefits of pretraining on reach decoder performance and training time, we evaluated the effect of pretraining the fUS-BMI on an example session (**Fig. 6E-G**). As with the saccade decoders, pretraining significantly shortened training time. In some cases, pretraining rescued a “bad” model. For example, the example session using only current data (**Fig. 6C**) displayed a heavy bias towards the left. When we used this same example session to pretrain the fUS-BMI a few days later, the new model made balanced predictions (**Fig. 6F**). The searchlight analysis for this example session revealed that the same dorsal LIP region from the example session without pretraining contained the majority of the most informative voxels (**Fig. 6G**). MIP and Area 5 also contained a few patches of highly informative voxels.

#### Using only data from the current session (**Fig. 7**-left)

The mean decoder accuracy reached significance by the end of each session (1 real-time and 3 simulated). The performance reached 65 ± 2% correct and took 67.67 ± 18.77 trials to reach significance.

**Figure 7.**
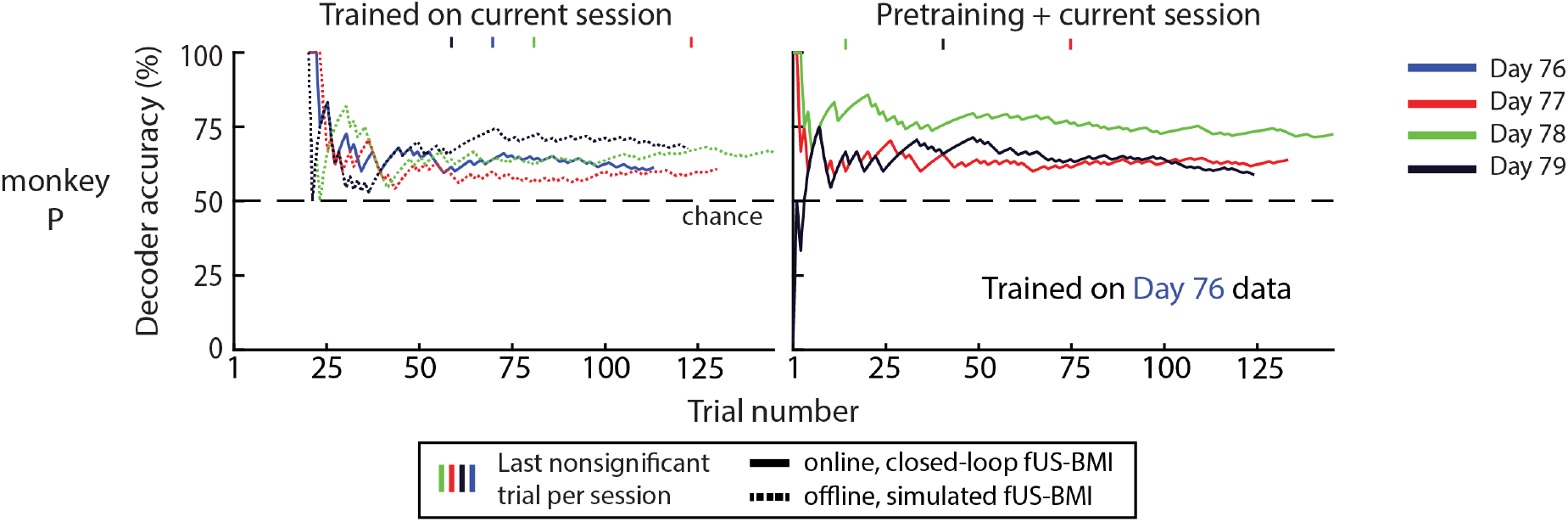
Performance across sessions for decoding two reach directions. Performance across sessions for monkey P. Same format as **Fig. 3A**.

#### Pretraining with data from a previous session (Fig. 7-right)

The mean decoder accuracy reached significance by the end of each session (3 real-time). Monkey P’s performance reached 65 ± 4% correct and took 43.67 ± 17.37 trials to reach significance. For two of the three real-time sessions, the number of trials needed to reach significance decreased with pretraining (-2 – 46 trials faster; 0-16 minutes faster). There was no statistical difference in performance between the sessions with and without pretraining (paired t-test, p<0.05). We also simulated the effects of not using any training data from the current session, i.e., using only the pretrained model (**Fig. S2C**). We did not observe a statistically significant difference between the performance (accuracy or number of trials to significant performance) whether current session training data was included or not.

These results are consistent with our previous study’s results^10^ that we can decode not only eye movements, but also reach movements. As with the eye movement decoders, we could pretrain the fUS-BMI using a previous session’s data and reduce, or even eliminate, the need for new training data.

## DISCUSSION

This work demonstrates, for the first time, a closed-loop, online, ultrasonic BMI and makes two other key advances that prepare for future work on the next generation of minimally-invasive ultrasonic BMIs: decoding more movement directions and stabilizing decoders across more than a month.

### Decoding more movement directions

We successfully decoded eight movement directions in real-time, an advance on previous work that decoded two saccade directions and two reach directions using pre-recorded data^10^. Specifically, we replicated the two direction results using real-time online data (**Fig. 2, 3, 6**, and **7**) and then extended the decoder to work for eight movement directions (**Fig. 4** and **5**).

### Stabilizing decoder across time

Electrode-based BMIs are particularly adept at sensing fast changing (∼10s of ms) neural activity from spatially localized regions (<1 cm) during behavior or stimulation that is correlated to activity in such spatially specific regions, e.g., M1 for motor and V1 for vision. Electrodes, struggle to track individual neurons over longer periods of time, e.g., between subsequent recording sessions. Consequently, decoders are typically retrained every day^18^. In this paper, we demonstrated a novel alignment method that stabilizes image-based BMIs across more than a month and decode from the same neural populations with minimal, if any, retraining. This is a critical development that enables easy alignment of previous days’ models to a new day’s data and begin decoding while acquiring minimal to no new training data. Much effort has focused on reducing or eliminating re-training in electrode-based BMIs^19–24^. These methods require identification of manifolds and/or latent dynamical parameters and collecting new data to align to these manifolds/parameters. Furthermore, some of the algorithms are computationally expensive and/or difficult to implement in online use. Our novel decoder alignment algorithm leverages the intrinsic spatial resolution and field of view provided by fUS neuroimaging to simplify this process in a way that is intuitive, repeatable, and performant.

### Improving performance

An ideal fUS-BMI would have performance that is better than the 45% correct at decoding eight movement directions found in this study. We have several ideas about how to improve the fUS-BMI performance.

*First*, realigning the recording chamber and ultrasound transducer along the intraparietal sulcus axis would allow sampling from a larger portion of LIP and MIP. In this paper, we placed the chamber and probe in a coronal orientation to aid anatomical interpretability. However, most of our imaging plane is not contributing to the decoder performance (**Fig. 2C, 2F, 4C, 4F, 6D, 6G**). Previous papers have found that receptive fields are anatomically organized along anterior-posterior and dorsal-ventral gradients within LIP^25^. By realigning the recording chamber orthogonal to the intraparietal sulcus in future studies, we could sample from a larger anterior-posterior portion of LIP with diverse range of directional tunings.

*Second*, we were limited to 2D planar imaging. The advent of 3D ultrafast volumetric imaging based on matrix or row-column array technologies will be capable of sensing changes in CBV from blood vessels that are currently orthogonal to the imaging plane. Additionally, 3D volumetric imaging can fully capture entire functional regions *and* sense multiple functional regions simultaneously. There are many regions which could fit inside a single 3D probe’s field of view and contribute to a motor BMI, for example: PPC, primary motor cortex (M1), dorsal premotor cortex (PMd), and supplementary motor area (SMA). These areas encode different aspects of movements including goals, sequences, and expected value of actions^26– 29^. This is just one example of myriad advanced BMI decoding strategies that will be made possible by synchronous data across brain regions. Currently, high-quality low-latency real-time 3D fUS imaging is not possible due to bandwidth, memory, and compute limitations. However, ongoing advances in hardware and algorithms will likely soon enable 3D fUS-BMI.

*Third*, another route for improved performance would be using more advanced decoder models. Convolutional neural networks are tailor-made for identifying image characteristics and are robust to spatial perturbation common in fUS images. Recurrent neural networks and transformers use “memory” processes that may be particularly adept at characterizing the temporal structure of fUS time series data. A potential downside of artificial neural networks (ANNs) like these is that they require significantly more training data. However, the methods presented here for across-session image alignment allow for pre-recorded data to be aggregated and organized into a large data corpus. Such a data corpus should be sufficient for training many ANNs. The amount of training data required could be further reduced by decreasing the feature count of the ANNs themselves. For example, one could reduce the input layer dimensions by restricting the data to features collected only from the task-relevant areas, such as LIP and MIP, instead of the entire image. ANNs additionally take longer to train (∼minutes instead of seconds) and would require different strategies for online retraining than used in this paper. In this paper, we leveraged the ability to rapidly retrain linear models on every trial without stopping the task. Although online training and inference using ANNs are beyond the scope of this paper, this is an active area of investigation for fUS-BMIs.

### Advantages of fUS neuroimaging

fUS neuroimaging has several advantages compared to existing BMI technologies. The large and deep field of view allows us to reliably record from multiple cortical and subcortical regions simultaneously – and to record from the same populations in a stable manner over long periods of time. fUS neuroimaging is epidural, i.e., does not require penetration of the dura mater, substantially decreasing surgical risk, infection risk, and tissue reactions while enabling chronic imaging over long periods of time (potentially many years) with minimal, if any degradation, in signal quality. In our NHP studies, fUS neuroimaging has been able to image through the dura, including the granulation tissue that forms above the dura (several ∼mm) with minimal loss in sensitivity.

In this paper, we demonstrated a new benefit of fUS: decoders that are stable across multiple days or even months. Using conventional image registration methods, we can align our decoder across different recording sessions and achieve excellent performance without collecting additional training data. A weakness of this current fUS-BMI compared to current electrophysiology BMIs is poorer temporal resolution. Electrophysiological BMIs have temporal resolutions in the 10s of milliseconds (e.g., binned spike counts). fUS can reach a similar temporal resolution (up to 500 Hz in this work) but is limited by the time constant of mesoscopic neurovascular coupling (∼seconds). Despite this neurovascular response acting as a low pass filter on each voxel’s signal, faster fUS acquisition rates can measure temporal variation across voxels down to <100 ms resolution. Earlier researchers already exploited this to track dynamic propagation of local hemodynamic changes through cortical layers and between functional regions within a single plane of view in NHPs^30^. As the temporal resolution and latency of real-time fUS imaging improves with enhanced hardware and software, tracking the propagation of these rapid hemodynamic signals may enable improved BMI performance and response time. Additionally, for the current study, and for many BMI applications, the goals of an action can be extracted despite the slow mesoscopic hemodynamic response and do not require the short latency required for extracting faster signals such as the trajectories of intended movements. Beyond movement, many other signals in the brain may be better suited to the spatial and temporal strengths of fUS, for example, monitoring biomarkers of neuropsychiatric disorders (discussed in detail below).

### Decoding hand, eye, or both?

Dorsal and ventral LIP contained the most informative voxels when decoding eye movements (**Fig. 2C, 2F, 4C, 4F**). This is consistent with previous literature that LIP is important for spatially specific oculomotor intention and attention^16,31,32^. Dorsal LIP, MIP, Area 5, and Area 7 contained the most informative voxels during reach movements (**Fig. 6D, 6G**). The voxels within the LIP closely match with the most informative voxels from the 2-direction saccade decoding, suggesting that our fUS-BMI may be using eye movement plans to build its model of movement direction. The patches of highly informative voxels within MIP and Area 5 suggest the fUS-BMI may also be using reach-specific information^14,33–36^. Future experiments will be critical for disentangling the mesoscopic contributions of LIP, MIP, Area 5, and other PPC regions for accurate effector predictions with a fUS-BMI. One such experiment would be recording and ultimately decoding fUS signals from the PPC as NHPs perform dissociated eye and reach movements^14^. As this fUS-BMI is translated into human applications, these effector-specific signals can also be more cleanly studied by instructing subjects to perform dissociated effector tasks.

### Moving beyond a motor BMI

The vast majority of BMIs have focused on motor applications, e.g. restoring lost motor function in people with paralysis^1^. Recently there has been interest in developing closed-loop BMIs to restore function to other demographics, such as patients disabled from neuropsychiatric disorders^37^. Depression, the most common neuropsychiatric disorder, affects an estimated 3.8% of people (280 million) worldwide^38^. In this paper, we demonstrated the utility of the fUS-BMI for motor applications to allow easier comparison with existing BMI technologies. fUS-BMIs seem to be an ideal platform for applications that require monitoring neural activity over large regions of the brain and long time scales, from hours to months. As demonstrated here, fUS neuroimaging captures neurovascular changes on the order of a second and is also stable over more than 1 month. Combined with neuromodulation techniques such as focused ultrasound, it may be possible to not only record from these distributed corticolimbic populations but also precisely modulate specific mesoscopic populations^39,40^.

The contributions presented here demonstrate the first online, closed-loop fUS-BMI. It prepares for the next generation of BMIs that are less invasive, high-resolution, stable across time, and scalable to sense activity from large regions of the brain. These advances are a step toward fUS-BMI for a broader range of applications, including restoring function to patients suffering from debilitating neuropsychiatric disorders.

## METHODS

### Experimental model and subject details

We implanted two healthy 14-year-old male rhesus macaque monkeys (*Macaca mulatta*) weighing 14-17 kg. All surgical and animal care procedures were approved by the California Institute of Technology Institutional Animal Care and Use Committee and complied with the Public Health Service Policy on the Humane Care and Use of Laboratory Animals.

### General

We used NeuroScan Live software (ART INSERM U1273 & Iconeus, Paris, France) interfaced with MATLAB 2019b (MathWorks, Natick, MA, USA) for the real-time fUS-BMI and MATLAB 2021a for all other analyses.

#### Animal preparation and implant

For each NHP, we placed a cranial implant containing a titanium head post and a craniotomy positioned over the posterior parietal cortex. The dura underneath the craniotomy was left intact. The craniotomy was covered by a 24 × 24 mm (inner dimension) chamber. For each recording session, we used a custom 3D-printed polyetherimide slotted chamber plug that held the ultrasound transducer. This allowed the same anatomical planes to be consistently acquired on different days.

#### Behavioral setup

NHPs sat in a primate chair facing a monitor or touchscreen. The LCD monitor was positioned ∼30 cm in front of the NHP. The touchscreen was positioned on each day so that the NHP could reach all the targets on the screen with his fingers but could not rest his palm on the screen. This was ∼20 cm in front of the NHP. Eye position was tracked at 500 Hz using an infrared eye tracker (EyeLink 1000, Ottawa, Canada). Touch was tracked using a touchscreen (Elo IntelliTouch, Milpitas, California). Visual stimuli were presented using custom Python 2.7 software based on PsychoPy^41^. Eye and hand position was recorded simultaneously with the stimulus and timing information and stored for offline analysis.

#### Behavioral tasks

NHPs performed several different memory-guided movement tasks. In the memory-guided saccade task, NHPs fixated on a center cue for 5 ± 1 seconds. A peripheral cue appeared for 400 ms in a peripheral location (either chosen from 2 or 8 possible target locations) at 20° eccentricity. The NHP kept fixation on the center cue through a memory period (5 ± 1 s) where the peripheral cue was not visible. The NHP then executed a saccade to the remembered location once the fixation cue was extinguished. If the NHP’s eye position was within a 7° radius of the peripheral target, the target was re-illuminated and stayed on for the duration of the hold period (1.5 ± 0.5 s). The NHP received a liquid reward of 1000 ms (0.75 mL) for successful task completion. There was an 8 ± 2 second intertrial interval before the next trial began. Fixation, memory, and hold periods were subject to timing jitter sampled from a uniform distribution to prevent the NHP from anticipating task state changes.

The memory-guided reach task was similar, but instead of fixation, the NHP used his fingers on a touchscreen. Due to space constraints, eye tracking was not used concurrently with the touchscreen, i.e., only hand or eye position was tracked, not both.

For the memory-guided BMI task, the NHP performed the same fixation steps using his eye or hand position, but the movement phase was controlled by the fUS-BMI. Critically, the NHP was trained to not make an eye or hand movements from the center cue until at least the reward was delivered. For this task variant, the NHP received a liquid reward of 1000 ms (0.75 mL) for successfully maintaining fixation/touch and correct fUS-BMI predictions. The NHP received a 100 ms (0.03 mL) reward for successfully maintaining fixation/touch during incorrect fUS-BMI predictions. This was done to maintain NHP motivation.

### fUS-BMI

#### Functional ultrasound sequence and recording

During each fUS-BMI session, we placed the ultrasound transducer (128-element miniaturized linear array probe, 15.6 MHz center frequency, 0.1 mm pitch, Vermon, France) on the dura with ultrasound gel as a coupling agent. We consistently positioned the ultrasound transducer across recording sessions using a slotted chamber plug. The imaging field of view was 12.8 mm (width) by 13-20 mm (height) and allowed the simultaneous imaging of multiple cortical regions, including lateral intraparietal area (LIP), medial intraparietal area (MIP), ventral intraparietal area (VIP), Area 7, and Area 5.

We used a programmable high-framerate ultrasound scanner (Vantage 256; Verasonics, Kirkland, WA) to drive the ultrasound transducer and collect pulse echo radiofrequency data. We used different plane-wave imaging sequences for real-time and anatomical fUS neuroimaging.

#### Real-time low-latency fUS neuroimaging

We used a custom-built computer running NeuroScan Live (ART INSERM U1273 & Iconeus, Paris, France) attached to the 256-channel Verasonics Vantage ultrasound scanner. This software implemented a custom plane-wave imaging sequence optimized to deliver Power Doppler images in real-time at 2 Hz with minimal latency between ultrasound pulses and Power Doppler image formation. The sequence used a pulse-repetition frequency of 5500 Hz and transmitted plane waves at 11 tilted angles equally spaced from -6° to 6°. These tilted plane waves were compounded at 500 Hz. Power Doppler images were formed from 200 compounded B-mode images (400 ms). To form the Power Doppler images, the software used an ultrafast Power Doppler sequence with an SVD clutter filter^42^ that discarded the first 30% of components. The resulting Power Doppler images were transferred to a MATLAB instance in real-time and used for the fUS-BMI. The prototype 2 Hz real-time fUS system had approximately an 800 ms latency from the end of the ultrasound pulse sequence to arrival of the beamformed fUS image in MATLAB. Each fUS image and associated timing information were saved for *post hoc* analyses.

#### Anatomical Doppler neuroimaging

At the start of each recording session, we used a custom plane-wave imaging sequence to acquire an anatomical image of the vasculature. We used a pulse repetition frequency of 7500 Hz and transmitted plane waves at 5 angles ([-6°, -3°, 0°, 3°, 6°]) with 3 accumulations. We coherently compounded these 5 angles from 3 accumulations (15 images) to create one high-contrast ultrasound image. Each high-contrast image was formed in 2 ms, i.e., at a 500 Hz framerate. We formed a Power Doppler Image of the NHP brain using 250 compounded B-mode images collected over 500 ms. We used singular value decomposition to implement a tissue clutter filter and separate blood cell motion from tissue motion^42^.

#### fUS-BMI Overview

There were three components to decoding movement intention in real-time: 1) apply preprocessing to a rolling data buffer, 2) train the classifier, and 3) decode movement intention in real-time using the trained classifier. As described previously^10^, the time for preprocessing, training, and decoding was dependent upon several factors, including the number of trials in the training set, CPU load from other applications, the field of view, and classifier algorithm (PCA+LDA vs cPCA+LDA). In the worst cases during offline testing, the preprocessing, training, and decoder respectively took approximately 10, 500 ms, and 60 ms. See ^10^ for more description of the amount of time needed for preprocessing, training, and prediction with different sized training sets.

#### Data preprocessing

Before feeding the Power Doppler images into the classification algorithm, we applied two preprocessing operations to a rolling 60-frame (30 seconds) buffer. We first performed a rolling voxel-wise z-score over the previous 60 frames (30 seconds) and then applied a pillbox spatial filter with a radius of 2 pixels to each of the 60 frames in the buffer.

#### Real-time classification

The fUS-BMI made a prediction at the end of the memory period using the preceding 1.5 seconds of data (3 frames) and passed this prediction to the behavioral control system via a TCP-based server (**Fig. S3**). We used different classification algorithms for fUS-BMI in the 2-direction and 8-direction tasks. For decoding two directions of eye or hand movements, we used class-wise principal component analysis (cPCA) and linear discriminant analysis (LDA), a method well suited to classification problems with high dimensional features and low numbers of samples^43,44^. This method is mathematically identical to that used previously for offline decoding of movement intention^10^ but has been optimized for online training and decoding. Briefly, we used cPCA to dimensionally reduce the data while keeping 95% variance of the data. We then used LDA to improve the class separability of the cPCA-transformed data. For more details on the method and implementation, see ^10,43,44^.

For decoding eight directions of eye movements, we used a multicoder approach where the horizontal (left, center, or right) and vertical components (down, center, or up) were separately predicted and combined to form the final prediction. As a result of this separate decoding of horizontal and vertical movement components, “center” predictions are possible (horizontal – center and vertical – center) despite although this not being one of the eight possible peripheral target locations. To perform the predictions, we used principal component analysis (PCA) and LDA. We used the PCA to reduce the dimensionality of the data while keeping 95% of the variance in the data. We then used LDA to predict the most likely direction. We opted for the PCA+LDA method over the cPCA+LDA for 8-direction decoding because we found in offline analyses that the PCA+LDA multicoder outperformed cPCA+LDA for decoding eight movement directions with a limited number of training trials.

#### Real-time training of model

We retrained the fUS-BMI classifier during the intertrial interval (without stopping the experiment) every time the training set was updated. For the real-time experiments, the data recorded during a successful trial was automatically added to the training set. During the initial training phase, successful trials were defined as the NHP performing the movement to the correct target and receiving his juice reward. Once in BMI mode, successful trials were defined as a correct prediction plus the NHP maintaining fixation until juice delivery.

For experiments that used a model trained on data from a previous session, we used data from all valid trials from the previous session upon initialization of the fUS-BMI. A valid trial was defined as any trial that reached the prediction phase, regardless of whether the correct class was predicted. The classifier then retrained after each successful trial was added to the training set during the current session.

#### Post hoc experiments

These experiments analyzed the effect of using only data from a single session on decoder performance. We simulated an online scenario where we trained and/or decoded on each attempted trial in order. We considered all trials where the monkey received a reward as successful and retrained after each trial.

#### *Connection with the behavioral system* – (**Fig. S3**)

We designed a threaded TCP server in Python 2.7 to receive, parse, and send information between the computer running the PsychoPy behavior software and the real-time fUS-BMI computer. Upon queries from the fUS-BMI computer, this server transferred task information, including task timing and actual movement direction, to the real-time ultrasound system. The client-server architecture was specifically designed to prevent data leaks: the actual movement direction was never transmitted to the fUS-BMI until after a successful trial had ended. The TCP server also received the fUS-BMI prediction and passed it to the PsychoPy software when queried. The average server write-read-parse time was 31 +/-1 (mean ± STD) ms during offline testing between two desktop computers (Windows) on a local area network (LAN).

### Across-session alignment

At the beginning of each experimental session, we acquired an anatomical image showing the vasculature within the imaging field of view. For sessions where we used previous data as the initial training set for the fUS-BMI, we then performed a semi-automated intensity-based rigid-body registration between the new anatomical image and the anatomical image acquired in a previous session. We used the MATLAB ‘imregtform’ function with the mean square error metric and a regular step gradient descent optimizer to generate an initial automated alignment of the previous anatomical image to the new anatomical image. If the automated alignment had misaligned the two images, the software prompted the proctor to manually shift and rotate the previous session’s anatomical image using a custom MATLAB GUI. We then applied the final rigid body transform to the training data from the previous session, thus aligning the previous session to the new session.

### Quantification and statistical analysis

Unless reported otherwise, summary statistics reported as XX ± XX are mean ± SEM.

#### Post hoc simulated session

We used the recorded real-time fUS images to simulate the effects of different parameters on fUS-BMI performance, such as using only current session data without pretraining. To do this, we streamed pre-recorded fUS images and behavioral data, frame by frame, to the same fUS-BMI function used for closed-loop, online fUS-BMI. To dynamically build the training set, we added all trials reaching the end of the memory phase regardless of whether the offline fUS-BMI predicted the correct movement direction. This was done because the high possible error rate from bad predictions meant that building the training set from only correctly predicted trials could lead to imbalanced samples across conditions (directions) and possibly contain insufficient trials to train the model; zero correct predictions for certain directions could prevent the model from ever predicting that direction.

#### Searchlight analysis

We defined a circular region of interest (ROI; 200 μm radius) and moved it sequentially across all voxels (left to right then top to bottom) in the imaging field of view. For each ROI, we performed offline decoding with 10-fold cross-validation using either the cPCA+LDA (2-directions) or PCA+LDA (8-directions) algorithm where we only used the voxels fully contained with each ROI. We assigned the mean performance across the cross-validation folds to the center voxel of the ROI. To visualize the results, we overlaid the performance (mean absolute angular error or accuracy) of the 10% most significant voxels on the anatomical vascular map from the session.

## ACKNOWLEDGEMENTS

We thank Kelsie Pejsa for assistance with animal care, surgeries, and training. We thank Claire Rabut and Lydia Lin for helpful discussions. We thank Krissta Passanante for her illustrations. W.S.G was supported by an NEI F30 (NEI F30 EY032799), the Josephine de Karman Fellowship, and the UCLA-Caltech MSTP (NIGMS T32 GM008042). S.L.N was supported by the Della Martin Foundation. This research was supported by the National Institute of Health BRAIN Initiative (grant 1R01NS123663-01 to R.A.A., M.G.S., and M.T.), the T&C Chen Brain-Machine Interface Center, and the Boswell Foundation (R.A.A.).

## Author Contributions

W.S.G., S.L.N., V.C., M.T., M.G.S., and R.A.A. conceived the study; S.L.N. established the fUS neuroimaging sequences and T.D., B.-F.O., F.S., and M.T. wrote the acquisition software for 2 Hz real-time fUS neuroimaging; W.S.G. and S.L.N. wrote the code for the fUS-BMI; W.S.G. trained the NHPs and acquired the data; W.S.G., S.L.N., and G.C performed the data processing and analysis; W.S.G. and S.L.N. drafted the manuscript with substantial contributions from M.G.S and R.A.A., and all authors edited and approved the final version of the manuscript. V.C., C.L., M.T., M.G.S., and R.A.A. supervised the research.

