## Supplemental Figures for "Decoding Motor Plans Using a Closed-Loop Ultrasonic Brain-Machine Interface"

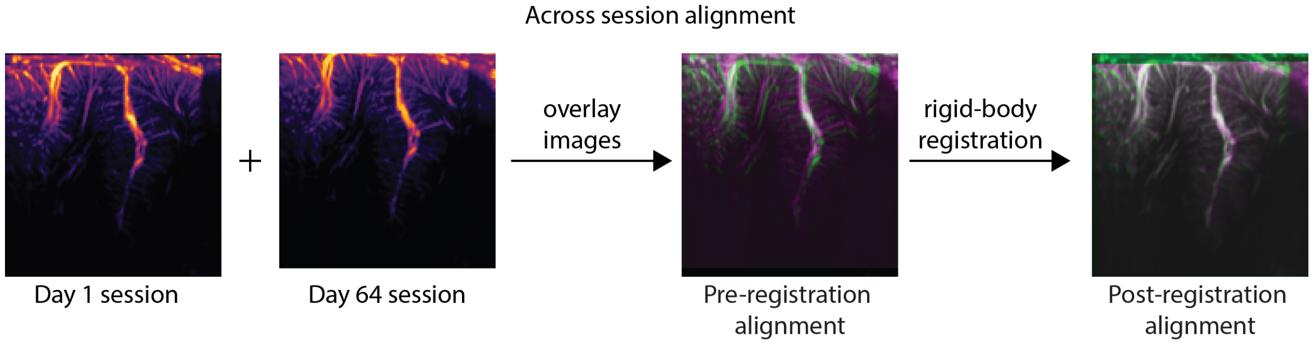


***Fig. S1 – Supplement to Fig. 1 – Across session alignment algorithm***
*We used semi-automated intensity-based rigid-body registration to find the transform from the previous session to the new imaging plane. The registration error is shown in the overlay where green represents the old session (Day 1) and magenta represents the new session (Day 64).*

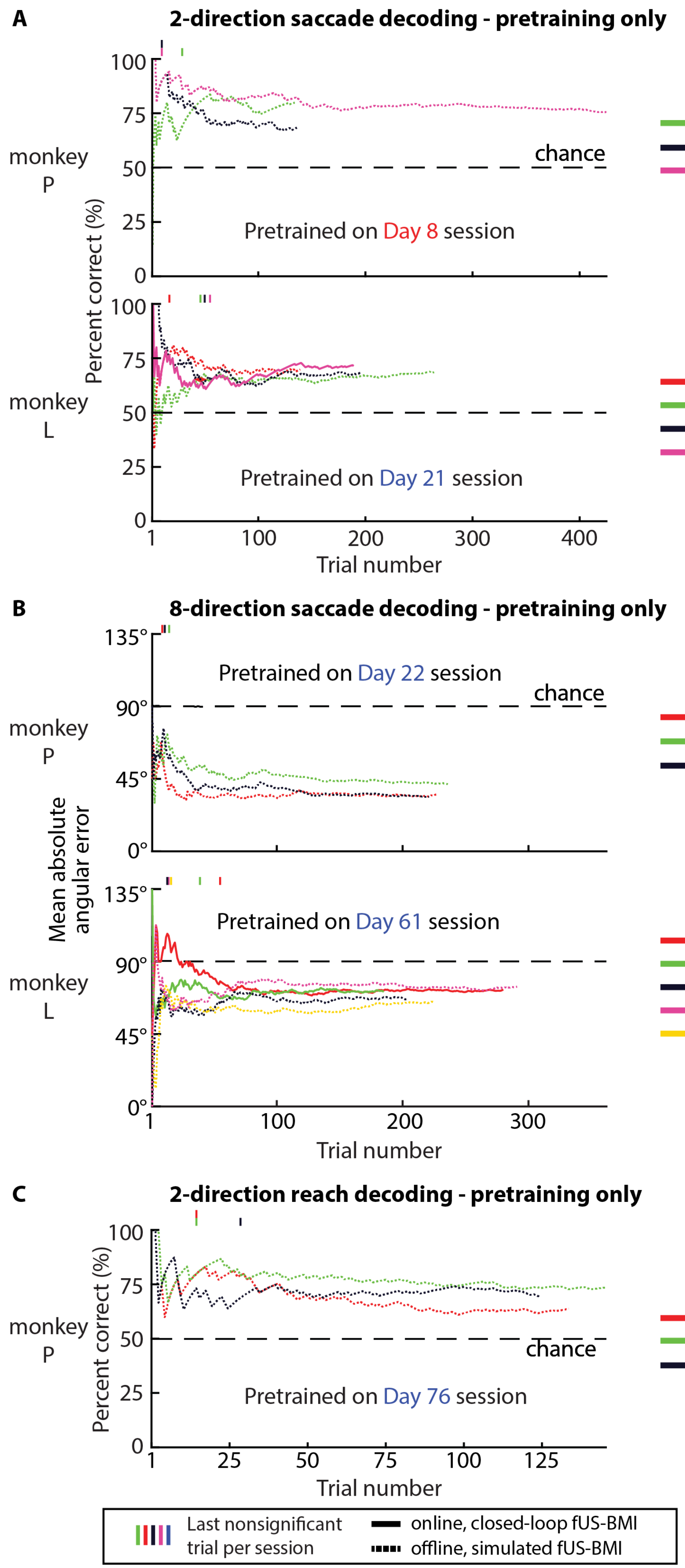


***Fig. S2 – Supplement to Fig. 3, 5, and 7 - Closed-loop, real-time decoding of movement directions using pretrained model only***

***A)*** *Performance for 2-direction saccade decoding using only the pretrained model. Same format as in **Fig. 3**.*

***B)*** *Performance for 8-direction saccade decoding using only the pretrained model. Same format as in **Fig. 5**.*

***C)*** *Performance for 2-direction reach decoding using only the pretrained model. Same format as in **Fig. 7**.*

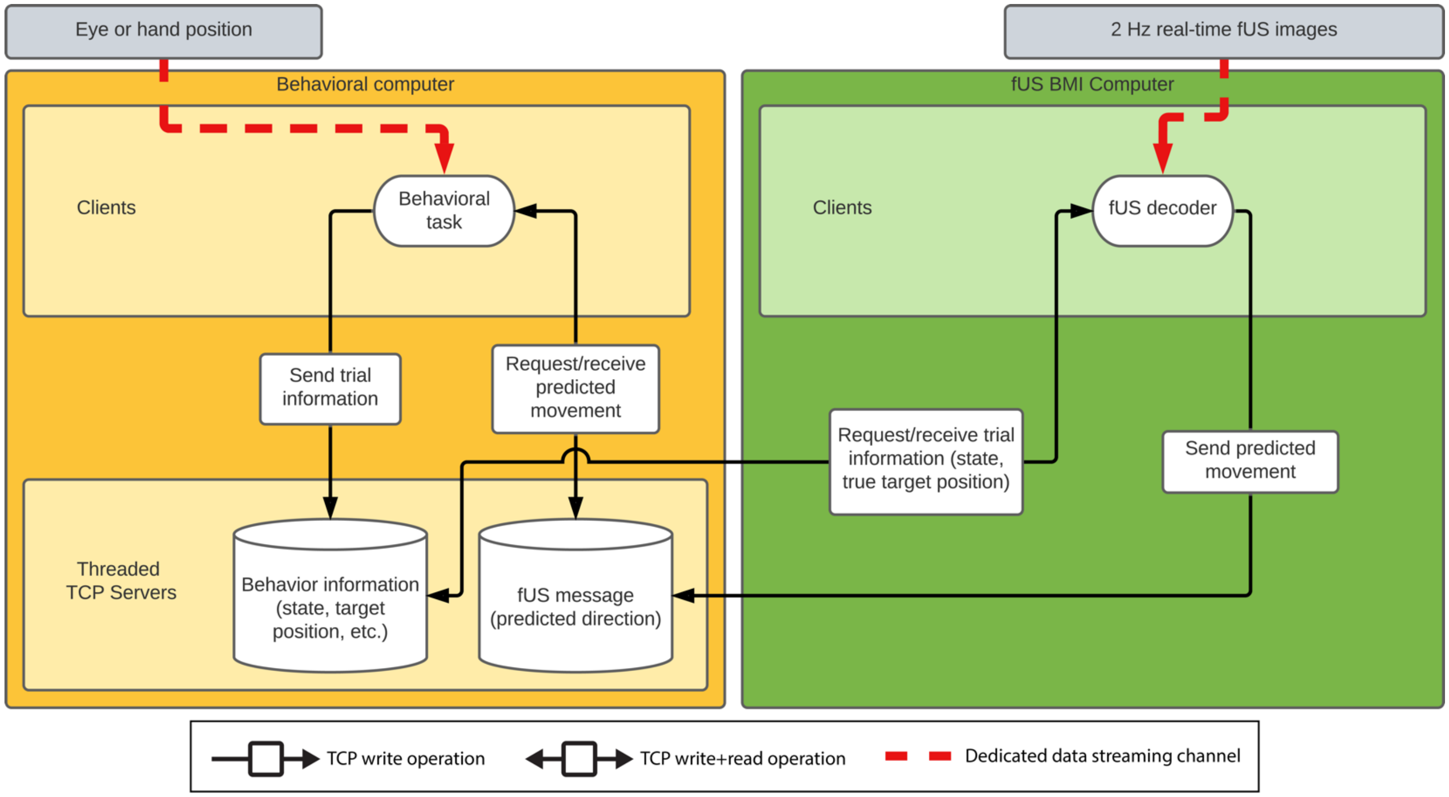


***Fig. S3 – TCP Communication Architecture for real-time fUS-BMI***

*We designed a threaded TCP server in Python 2.7 to receive, parse, and send information between the computer running the PsychoPy behavior software and the real-time fUS-BMI computer. Upon queries from the fUS-BMI computer ("fUS decoder"), this server transferred task information, including task timing and actual movement direction, to a real-time ultrasound system. The client-server architecture was specifically designed to prevent data leaks, i.e., the actual movement direction was never transmitted to the fUS-BMI until after a successful trial had ended. The TCP server also received the fUS-BMI prediction and passed it to the PsychoPy software when queried. The average server write-read-parse time was 31 +/- 1 (mean ± STD) ms during offline testing between two Windows computers on a local area network.*
